# A two-level, dynamic fitness landscape of hepatitis C virus revealed by self-organized haplotype maps

**DOI:** 10.1101/2021.04.22.441053

**Authors:** Soledad Delgado, Celia Perales, Carlos García-Crespo, María Eugenia Soria, Isabel Gallego, Ana Isabel de Ávila, Brenda Martínez-González, Lucía Vázquez-Sirvent, Cecilio López-Galíndez, Federico Morán, Esteban Domingo

**Author notes:** Both authors have contributed equally to this work; the names are written in alphabetical order (Soledad Delgado, Celia Perales). Addresses for correspondence.

## Abstract

Fitness landscapes reflect the adaptive potential of viruses. There is no information on how fitness peaks evolve when a virus replicates extensively in a controlled cell culture environment. Here we report the construction of Self-Organized Maps (SOMs), based on deep sequencing reads of three amplicons of the NS5A-NS5B-coding region of hepatitis C virus (HCV). A two-dimensional neural network was constructed and organized according to sequence relatedness. The third dimension of the fitness profile was given by the haplotype frequencies at each neuron. Fitness maps were derived for 44 HCV populations that share a common ancestor that was passaged up to 210 times in human hepatoma Huh-7.5 cells. As the virus increased its adaptation to the cells, the number of fitness peaks expanded, and their distribution shifted in sequence space. The landscape consisted of an extended basal platform, and a lower number of protruding higher fitness peaks. The function that relates fitness level and peak abundance corresponds a power law, a relationship observed with other complex natural phenomena. The dense basal platform may serve as spring-board to attain high fitness peaks. The study documents a highly dynamic, double-layer fitness landscape of HCV when evolving in a monotonous cell culture environment. This information may help interpreting HCV fitness landscapes in complex in vivo environments.

**IMPORTANCE:** The study provides for the first time the fitness landscape of a virus in the course of its adaptation to a cell culture environment, in absence of external selective constraints. The deep sequencing-based self-organized maps document a two-layer fitness distribution with an ample basal platform, and a lower number of protruding, high fitness peaks. This landscape structure offers potential benefits for virus resilience to mutational inputs.

## INTRODUCTION

High viral mutation rates lead to generation of complex and dynamic mutant spectra termed viral quasispecies, which are important for adaptability to changing environments (1). In the case of hepatitis C virus (HCV), quasispecies complexity in infected patients (quantified as the number of different genomes estimated to be present in the replicating mutant ensembles, as sampled from serum and liver samples) can exert an influence on disease progression and response to antiviral treatment [(2-4); reviewed in (5, 6)]. An understanding of the mechanisms that modify mutant distributions *in vivo* can be facilitated by minimizing the number of selective constraints during viral replication. This can be approached with cell culture systems that sustain long-term virus replication, as is the case of HCV replicating in Huh-7.5 cells (7-10).

The objective of the present work has been to determine fitness landscapes of sequential HCV populations replicating in a non-coevolving cellular environment, devoid of externally applied selective constraints. Only perturbations inherent to the cell culture and the changing mutant spectrum of the replicating virus were present (11). The study analyzes the haplotype relatedness and frequencies in a clonal HCV population, and its derivatives resulting from up to 210 serial passages in human hepatoma Huh-7.5 cells (equivalent to about 730 days of continuous replication) (11-14). The starting, clonal HCV population was generated by transcription of plasmid Jc1FLAG2(p7-nsGluc2A) (15), followed by RNA electroporation into Huh-Lunet cells, and minimum amplification of the progeny virus in Huh-7.5 cells (12). In this experimental design, fresh cells were infected with the virus shed into the cell culture medium of the previous infection, so that cellular evolution was prevented. Each passage involved infection of4×10^5^ Huh-7.5 reporter cells with 4 x 10^4^ to 4 x 10^6^ HCV TCID_50_ units (depending on the passage number) (11, 13). The multiplicity of infection (MOI) was 0.1 to 10 TCID_50_/cell. Under these conditions, possible distorting effects of stochasticity on quasispecies structure should be limited (16), and accumulation of defective genomes was largely avoided. This was suggested by the constancy of specific infectivity along 200 passages, and the fact that biological and molecular clones retrieved from the passaged virus displayed similar sequence diversification (11).

Our previous comparative analyses of the initial HCV population (termed HCV p0) and the populations at passages 100 and 200 (termed HCV p100 and HCV p200, respectively) revealed several phenotypic modifications, concomitantly with the process of adaptation to cell culture. The modifications included enhanced resistance to antiviral agents in the absence of specific inhibitor-escape amino acid substitutions (12, 17-21), as well as increases in virus particle density, in capacity to kill host cells, and in the extent of shutoff of host cell protein synthesis (13). Both, HCV p100 and HCV p200 exhibited a 2.3-fold increase in replicative fitness, relative to the initial population HCV p0 arbitrarily assigned a fitness of 1.0, as measured by growth-competition experiments in Huh-7.5 cells (13, 17). The reason why fitness did not increase from passage 100 to 200 may lie in viral population size limitations, as previously documented with vesicular stomatitis virus (22, 23). However, not all replicative parameters -calculated for each population individually—plateaued at passage 100. In a five serial passage test in Huh-7.5 cells, the intracellular exponential growth rate was 17- and 45-fold larger for HCV p100 and HCV p200, respectively, than for HCV p0 (13). In contrast, the maximum extracellular infectious progeny attained was 1.17-fold higher for both HCV p100 and HCV p200 than for HCV p0, in agreement with the leveling-off of fitness values denoted by the competition experiments (13).

Deep-sequencing of the genome of populations that were sampled to monitor the evolution from HCV p0 to HCV p200 revealed a large number of mutations that varied in frequency even between successive passages; we referred to the effect of these types of mutations as mutational waves (13). Strikingly, the waves did not subside when the population had increased its adaptation to the cellular environment since they were even more pronounced at late than at early viral passages (11, 24, 25). This seemingly paradoxical observation begged for an examination of the possible modifications underwent by the fitness landscape of individual HCV populations in their transition from HCV p0 to HCV p200.

Previous studies have afforded evidence that fitness landscapes for RNA viruses replicating in their natural environments are rugged and variable (16, 26-33). Fitness effects of mutations or amino acid substitutions have often been inferred from predicted or experimentally verified activity or stability of virus-coded proteins, or from the replicative performance of reconstructed viruses (34-39). An alternative approach has been to derive fitness landscapes from mutation frequencies calculated either from standard (consensus) sequences or from deep-sequencing data (27, 31, 38, 40).

There is no information on fitness landscapes of viral populations that have been extensively passaged in a cell culture environment, in absence of external selective constraints, as is the case of the evolution from HCV p0 to HCV p200. The abundance of genome types in a mutant spectrum is ranked according to relative fitness [reviewed in (16)]. This has been the rationale to investigate the fitness landscape of individual HCV populations, based on haplotype abundance derived from ultra-deep sequencing (UDS) reads. To this aim, we have applied an Artificial Neural Network (ANN) procedure as a learning method to derive Self-Organized Maps (SOM) (41, 42). The Kohonen’s SOM algorithm classifies a set of input data-vectors (in our case viral genomic sequences) in a bi-dimensional map. By an unsupervised process, it groups data vectors by similarity, projecting those vectors that have similar content in neighboring regions of the map (two-dimensional grid). In the case of viral genome sequences, the SOM algorithm generates an ordered grid in which each node (neuron) is associated with a reference RNA sequence (30, 43). Each neuron of the network maps all input sequences that fall within a distance from its reference vector which is smaller than the distance to the rest of reference vectors. Since vectors represent viral genomic sequences, to calculate numerical distances between vectors, a codification algorithm has been used, as previously described (43) (details are given in Materials and Methods and Fig. S1 in https://saco.csic.es/index.php/s/7TgiQcCr9ifpnt5). The SOM analysis of 44 HCV populations derived from HCV p0, HCV p100 and HCV p200 has disentangled the fitness landscape during long-term adaptation of HCV to Huh-7.5 cells. The SOM display has defined a remarkable HCV fitness topology consisting of a discrete number of high fitness peaks emerging from a lower fitness layer that approximates a fitness platform. The landscape is highly dynamic as evidenced by an almost complete shift in the region of sequence space occupied by the analyzed amplicons during the last 100 serial passages. Implications of the two-level fitness topology are discussed.

## RESULTS

### Self-organized maps and fitness landscape of mutant spectra of HCV populations, obtained from haplotype abundances

A clonal HCV p0 population derived from plasmid Jc1FLAG2(p7-nsGluc2A) (12, 15) was subjected to 200 serial passages in Huh-7.5 reporter cells, and samples from the initial population and from HCV p100 and HCV p200 were further passaged up to ten times in two separate experiments and several replicas, thus providing a total of 44 HCV populations for deep sequencing analysis (Fig. 1A). Three amplicons (termed A1, A2 and A3), extending from HCV genome residues 7649 to 8653 (residue numbering according to isolate JFH-1; accession number #AB047639) were analyzed (Fig. 1B). Amplicon A1 spans residues 7649 to 7960 (that correspond to amino acids 461 of NS5A to amino acid 98 of NS5B). A2 spans residues 7940 to 8257 (amino acids 92 to 197 of NS5B), and A3 covers residues 8231 to 8653 (amino acids 189 to 329 of NS5B). The number of processed, clean reads and the deduced number of haplotypes (number of identical reads represented by a nucleotide sequence) for the three amplicons are given in Table 1. For each of the 44 viral populations and amplicon, a FASTA file with the haplotype sequences, including the HCV genomic sequence contained in plasmid Jc1FLAG2(p7-nsGluc2A) —which is also used as reference for mutation counting—was prepared (Fig. S2 in https://saco.csic.es/index.php/s/7TgiQcCr9ifpnt5). The sequence of each haplotype was labeled with a name, the number of identical sequences that define it, and its frequency in each population (Fig. S3 in https://saco.csic.es/index.php/s/7TgiQcCr9ifpnt5). We employed the 3D irregular codification (43) to transform nucleotide sequences into numerical vectors. In this procedure, each nucleotide is located at a vertex of an irregular tetrahedron with distance 1 between A-G and C-U vertices, and distance 2 between the rest of pairs; that is, a distinction is made among mutation types. The codified sequences in each amplicon were used to train a SOM that comprised a set of neurons, each with a prototype vector, organized in a 15×15 two-dimensional (2D) neuron grid; a different SOM was trained for each amplicon. In this way, training sequences were mapped around the neuron with the prototype vector that best matched in terms of Euclidean distance [the “best matching” unit or bmu (41, 42); see Materials and Methods for additional information on SOM derivation)]. In our sequence analysis and processing, no insertion-deletions (*indels*) were recorded. [Their exclusion is justified by our evidence that they may arise artifactually in homopolymeric tracts upon RNA amplification (13)]. On the 2D neuron grid, the third dimension is given by the frequency of each group of sequences mapped around each neuron, thereby unfolding into a three-dimensional (3D) fitness map for the 44 HCV populations depicted in Fig. 1A. The 15 x 15 2D grids with sequence identification of each neuron, 3D maps, and tabulated numerical values for each population, experiment and amplicon are compiled in Figs. S4 to S6 in https://saco.csic.es/index.php/s/7TgiQcCr9ifpnt5, with additional numerical information in links quoted therein). Since no major differences were observed between experiment 1 and 2 and among parallel passage replicas, composite fitness maps that included all the populations derived either from HCV p0, HCV p100 or HCV p200, were obtained for each amplicon. The resulting maps (Fig. 2) reveal an expansion of the total number of haplotypes and fitness peaks in the evolution from HCV p0 to either HCV p100 or HCV p200 (fold-increase range of 2.1 to 5.3 for haplotypes, and of 2.0 to 4.0 for fitness peaks). Concerning the number of different haplotypes (given inside each panel of Fig. 2), the increase was significant in the evolution from HCV p0 to HCV p100 (p = 0.0439; t-test), and from HCV p0 to HCV p200 (p = 0.0015; t-test). The difference between HCV p100 and HCV p200 did not reach statistical significance (p = 0.2687; t-test).

**FIG 1.**
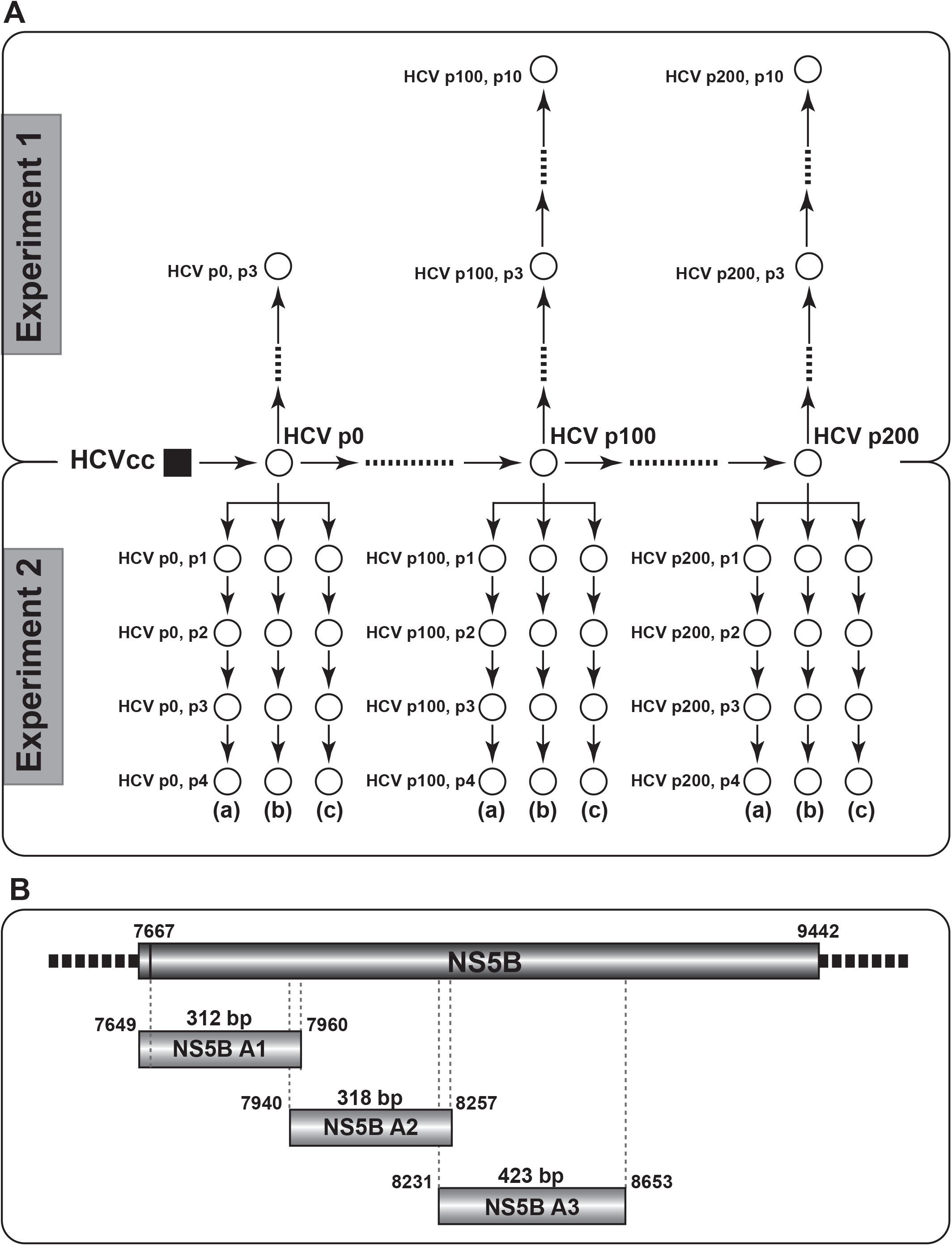
Experimental design and HCV amplicon analysis. (A) Schematic representation of the passages underwent by HCV p0 [derived from HCVcc (12); Materials and Methods)] in Huh-7.5 reporter cells. Populations are depicted as empty circles and passage number is indicated by p (HCV p100, p3 means population HCV p100 subjected to three passages in Huh-7.5 cells). Experiment 1 (upper part) and experiment 2 (lower part) were performed starting with samples of the same HCV p0, HCV p100 and HCV p200 populations. In experiment 2, (a), (b) and (c) indicate triplicate passage series carried out in parallel. A total of 44 HCV populations (corresponding to the empty circles) were analyzed by deep sequencing. The mutations (and deduced amino acid substitutions) identified in the populations from experiment 1 were reported in (18), and those in the populations from experiment 2 in (11). (B) HCV genomic residues 8261 (NS5A-coding region) to 9265 (NS5B-coding region) (genome numbering according to reference isolate JFH-1), and length in base pairs (bp) of amplicons A1, A2 and A3 analyzed by Illumina MiSeq sequencing. Note that the 21 most 3’terminal nucleotides of A1 are redundant with the 21 most 5’terminal nucleotides of A2, and that the 27 most 3’terminal nucleotides of A2 are redundant with the most 5’terminal nucleotides of A3. Further details on virus origin, GenBank accession numbers, and sequencing procedures are given in Materials and Methods.

**Table 1.**
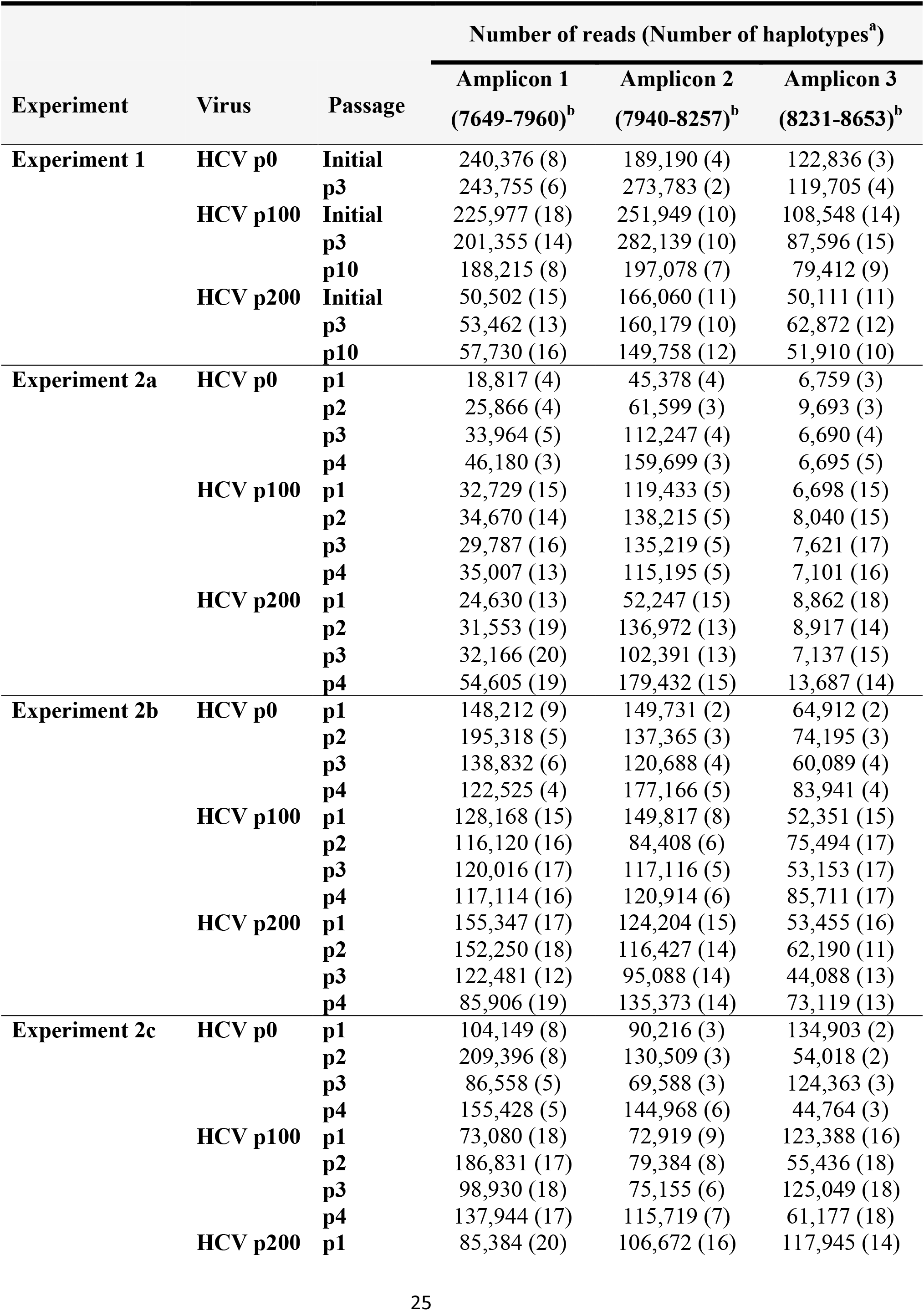
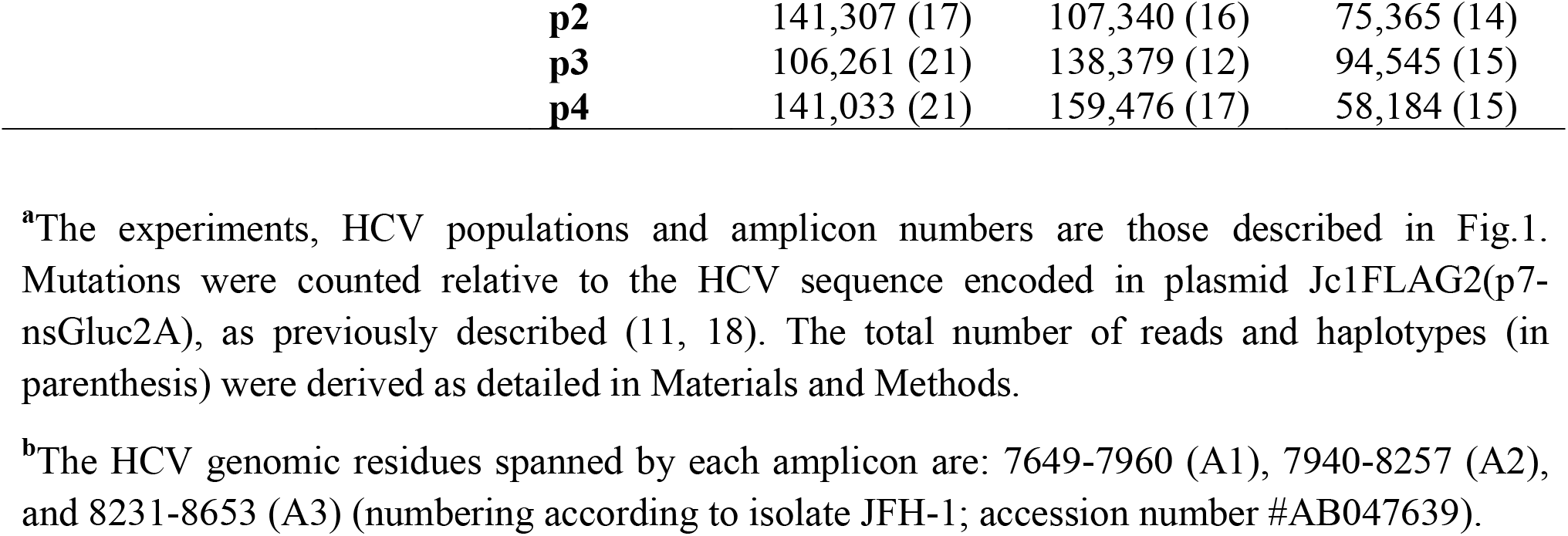
Number of reads and haplotypes derived from MiSeq Illumina sequencing of HCV amplicons A1, A2 and A3.

**FIG 2.**
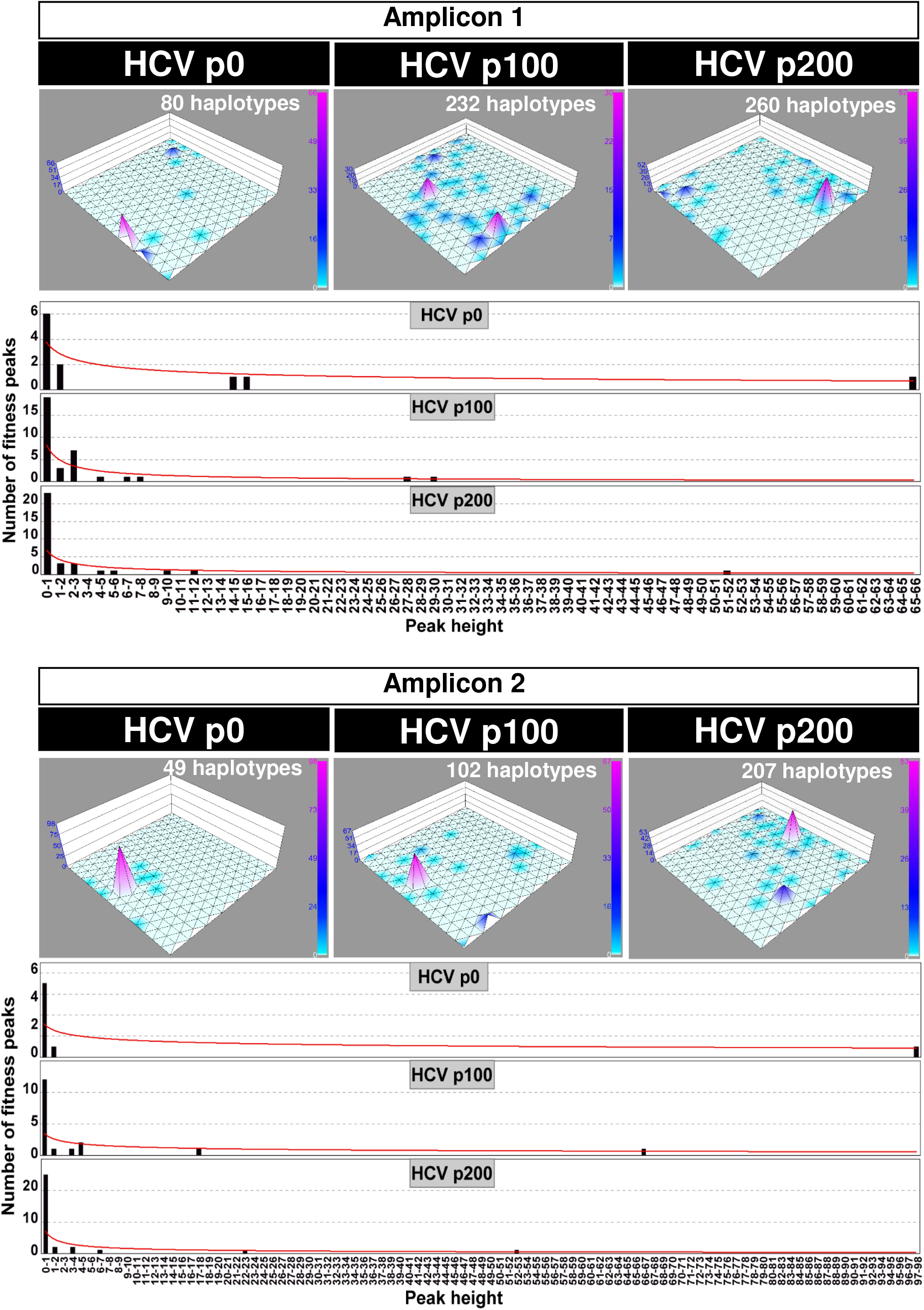
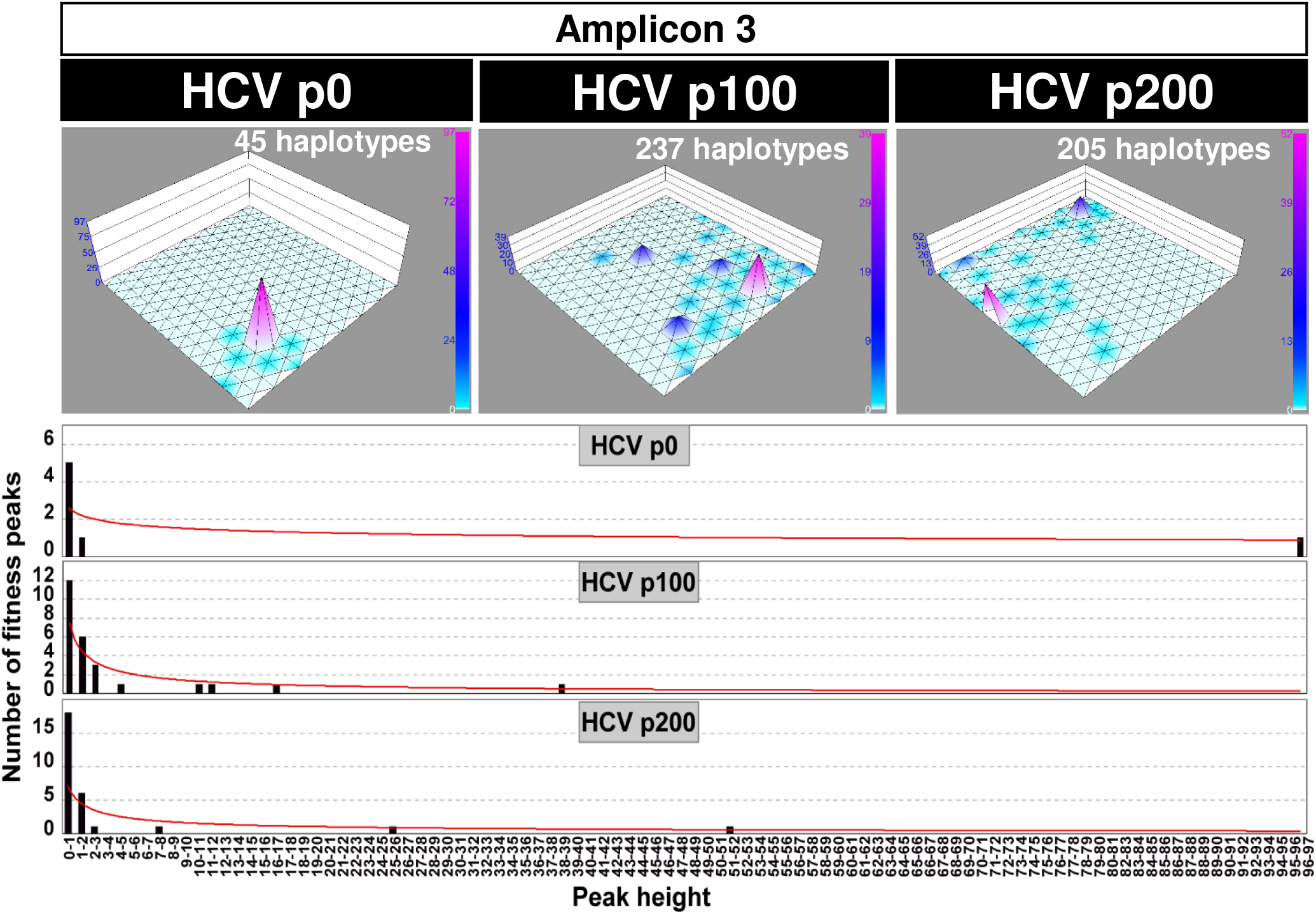
SOM-derived fitness maps, and number of fitness peaks distributed according to haplotype abundance. The amplicon number is indicated at the top of each panel and graphics group. The three 15 x 15 neuron grids for all populations derived either from HCV p0, HCV p100, or HCV p200 (displayed in Fig. 1A), with the total number of haplotypes that entered the analysis are indicated in each panel. Peak height is determined by sequence abundance, which is color coded with a scale included at the right of each fitness graph. The distribution of number of fitness peaks (ordinate) versus peak height (sequence abundance in unit range displayed in abscissa) is described by the following functions: Amplicon 1: HCV p0: y=3.7934x^-0.403^ (R^2^=0.7672); HCV p100: y=8.2657x^-0.77^ (R^2^=0.6463); HCV p200: y=6.6996x^-0.709^ (R^2^=0.5974). Amplicon 2: HCV p0: y=3.1334x^-0.281^ (R^2^=0.3804); HCV p100: y=3.4527x^-0.395^ (R^2^=0.3649); HCV p200: y=7.0358x^-0.638^ (R^2^=0.5818). Amplicon 3: HCV p0: y=2.5728x^-0.233^ (R^2^=0.3807); HCV p100: y=7.453x^-0.723^ (R^2^=0.7755); HCV p200: y=6.97x^-0.629^ (R^2^=0.5886). Note that scales are not the same in different panels. The origin of the sequences, derived haplotypes, and procedures are described in Materials and Methods.

The difference in the number of fitness peaks between HCV p0 and HCV p100 and between HCV p0 and HCV p200 was statistically significant (p = 0.0295 and p = 0.0004, respectively; t-test). The difference between HCV p100 and HCV p200 did not reach statistical significance (p = 0.1857; t-test). The distribution of the number of peaks as a function of peak height was indistinguishable for the three viral populations and amplicons (p = 1; chi-square test). In all cases, there is an accumulation of the number of fitness peaks within the peak height range 0-1 (graphics in Fig. 2). Considering the three amplicons together, the number of fitness peaks in range 0-1, 1-2, and all other range values was 18, 4, 5, respectively, for HCV p0; the corresponding values were 43, 10, 25 for HCV p100, and 66, 11, 17 for HCV p200 (data in Fig. 2). The bias is also evidenced by the ratio of fitness peaks with the minimal range (0-1) sequence frequency (the third dimension of the fitness maps color coded in Fig. 2) relative to the number of peaks that fall into any other range. The average ratio for all amplicons and populations was 0.66 (range 0.46-0.78). The dominance is also recapitulated in the function that relates the number of peaks with their sequence abundance at each neuron (third dimension in the fitness plot) (equations given in the legend for Fig. 2). Interestingly, the functions identify a power law for each virus and amplicon, unveiling for fitness maps a type of relationship found with other complex phenomena in physics and biology (see Discussion).

A shift in the occupation of sequence space (position of peaks in the 2D grid) was observed in all cases (Fig. 2). The position of the fitness peaks moved in their location in the three populations, except for the most prominent peak of amplicon 2 in HCV p0 that was also present in HCV p100. This shared peak was represented by a haplotype of identical sequence in each of the HCV p0 and HCV p100 populations that were integrated into the maps depicted in Fig. 2; the sequence was coincident with that present in the parental plasmid Jc1FLAG2(p7-nsGluc2A) (Table S1 in https://saco.csic.es/index.php/s/7TgiQcCr9ifpnt5); haplotype alignments are available in (https://saco.csic.es/index.php/s/586L2f9jJQtbRXq). The consistency of peak display among replicas of the same population (data given in Figs. S4 to S6 in https://saco.csic.es/index.php/s/7TgiQcCr9ifpnt5) validates the differences observed among different populations. Therefore, the replicative fitness increase in the evolution from HCV p0 to either HCV p100 or HCV p200 was reflected mainly in the number of low fitness peaks that occupied an increased, albeit shifting, portion of sequence space.

### Shared and unique fitness peaks among HCV populations

To express quantitatively the spread of mutant spectra in sequence space upon evolution from HCV p0 to HCV p100 and HCV p200, the number of shared and unique fitness peaks was recorded (Fig. 3). The ratio of number of unique peaks in HCV p0 relative to the number of peaks shared by the three populations was 2.5, 6 and 2 for amplicons A1, A2 and A3, respectively; the corresponding ratios were 14, 14, 11 for HCV p100, and 14, 30, 12.5 for HCV p200. The values were similar when the ratio was calculated relative to the number of peaks shared with any of the other populations; in this case, the ratios for HCV p0 were 2.5, 3, 4 for amplicons A1, A2, A3, respectively, and increased to 14, 7, 22 for HCV p100, and to 14, 30, 25 for HCV p200. For HCV p0, the difference between the number of unique versus shared peaks was not statistically significant: p = 0.50, p = 0.17, and p = 0.50 for amplicons A1, A2, and A3, respectively (proportion test). In contrast, for HCV p100 and HCV p200, the bias in favor of unique versus shared peaks was highly significant. For HCV p100 the p values obtained were p = 1.76 x 10^−7^, p = 0.00135, and p = 1.209 x 10^−6^ for amplicons A1, A2, and A3, respectively (proportion test). For HCV p200 the p values obtained were p = 1.76 x 10^−7^, p = 7.392 x 10^−12^, and p = 9.972 x 10^−9^ for amplicons A1, A2, and A3, respectively (proportion test). Therefore, the diversification and progressive occupation of sequence space by clonal HCV upon replication in Huh-7.5 cells is confirmed by the number of unique fitness maxima in HCV p100 and HCV p200. The largest increase was scored by amplicon 2, in agreement with the quantification of haplotypes and fitness peaks (compare Figs. 2 and 3).

**FIG 3.**
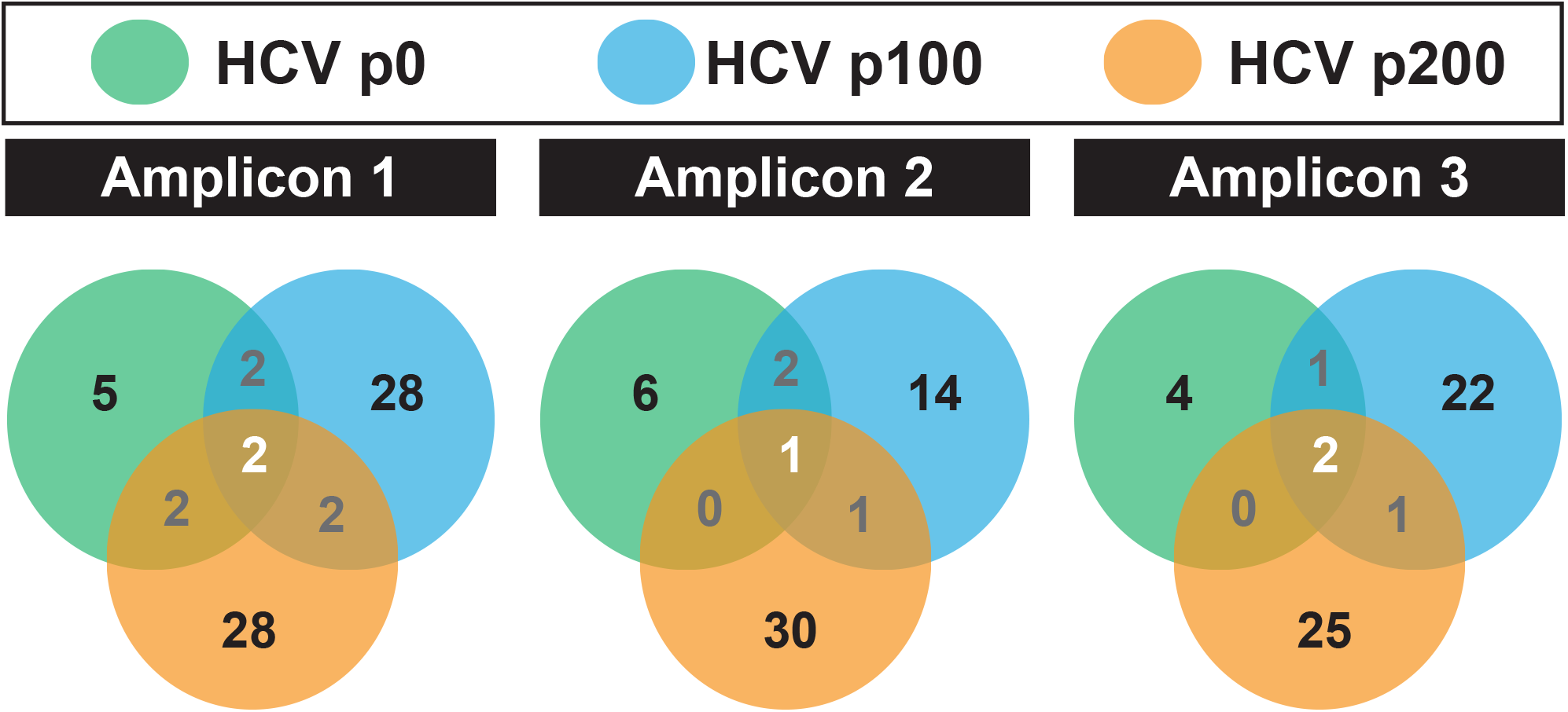
Distribution of fitness peaks among HCV populations. Venn diagrams indicating for each amplicon the number of peaks unique to one HCV population and those shared by two or more HCV populations. Populations are color coded. Peak identity was determined according to data summarized in Table 1, Fig. 2, and Supplemental Material (https://saco.csic.es/index.php/s/7TgiQcCr9ifpnt5).

### Fused amplicons

To produce a global image of the fitness landscape of the genomic region analyzed by incorporating the information of the three amplicons in a single graphic, it was necessary to equalize their length in nucleotides. Since the amplicons have overlapping sequences (Fig. 1B), we completed for each amplicon a length of 1005 nucleotides using the missing information provided by the other amplicons from the same population (procedure detailed in Materials and Methods). Then a 25 x 25 Kohonen’s ANN was trained using all the fused haplotypes. Fitness maps were built for each of the 44 populations, based on haplotype frequencies mapped around each neural unit (Figs. S7 to S9 in https://saco.csic.es/index.php/s/7TgiQcCr9ifpnt5). Since no major differences were noted among the individual fitness maps, a composite landscape was recapitulated for HCV p0, HCV p100 and HCV p200, each together with its derived populations. The results (Fig. 4) illustrate the peak dispersion upon evolution from HCV p0 to HCV p100 and HCV p200, and renders evident a striking location displacement of fitness peak abundance within the 2D grid. In particular, peaks in HCV p100 and HCV p200 clumped at opposite grid localities. Interestingly, in the process of amplicon fusion the power law that related number of fitness peaks with peak height for individual amplicons was no longer found for HCV p100 and HCV p200 (equations given in the legend for Fig. 4) (see Discussion). In conclusion, despite prolonged replication in a non-evolving cellular environment the HCV fitness landscape appears as remarkably broad, rugged, dynamic, and that approximates a two-layer peak height distribution.

**FIG 4.**
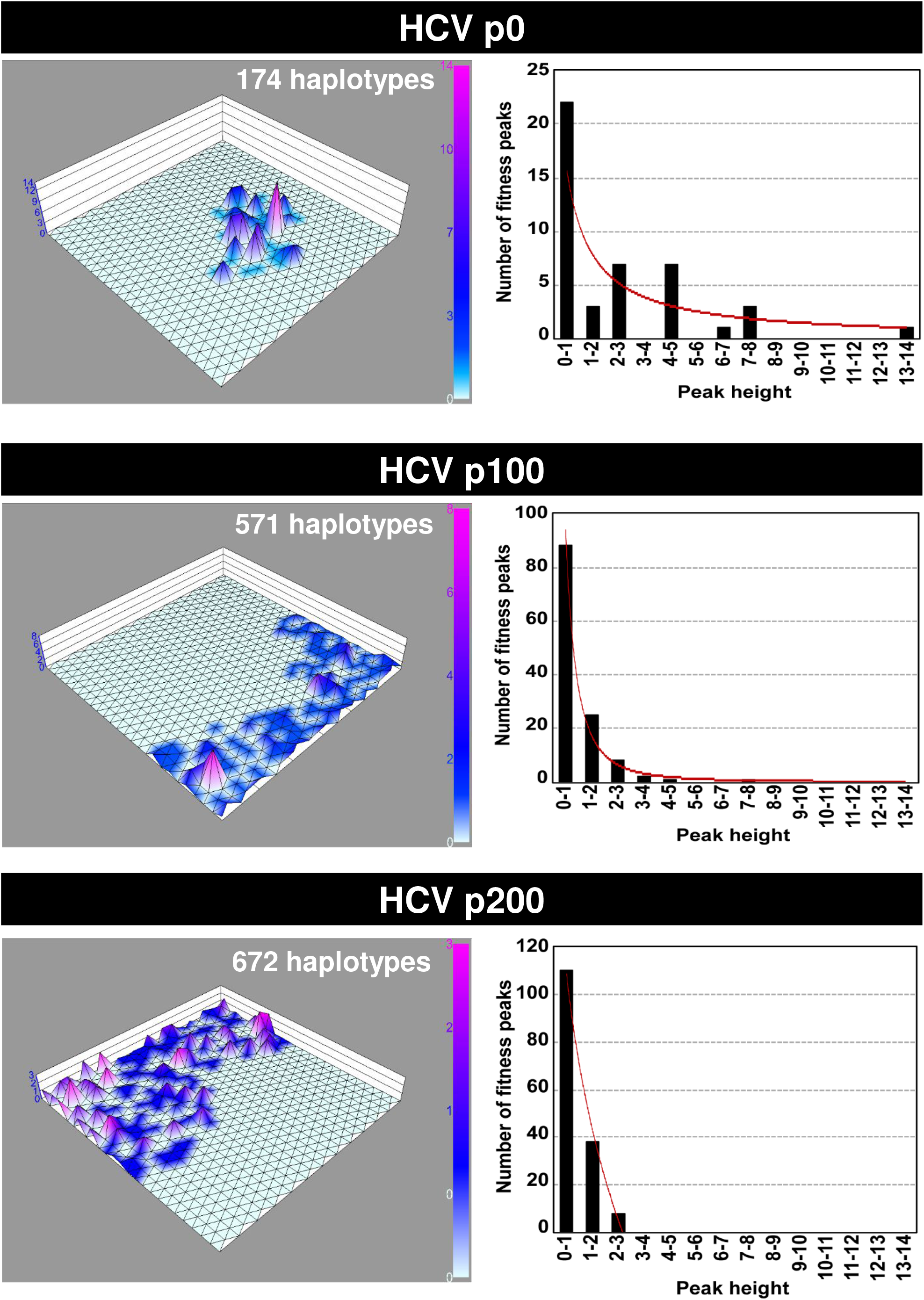
Fitness maps constructed with the fused NS5B amplicons. The HCV population and number of haplotypes used for the 25×25 neuron graphic are indicated on the left of each fitness map. Peak height is determined by sequence abundance, which is color coded with a scale included at the right of each map. The distribution of number of fitness peaks (ordinate) versus peak height (sequence abundance in unit range displayed in abscissa) is described by the following functions: HCV p0: y=15.659x^-1.001^ (R^2^=0.6519). HCV p100: y=93.588 x^2.441^ (R^2^=0.9931). HCV p200: -94.031ln(x) + 108.16 (R^2^=0.9931). Note that scales are not the same in different panels. Procedures are described in Materials and Methods.

### Mutation types and amino acid substitution tolerance in haplotypes from low and high fitness peaks

To analyze a possible difference in mutation types and amino acid substitution tolerance between sequences found in low and high fitness peaks, the peaks were divided in two groups: one with the sequences that populate fitness peaks of height range 0-1, and another group with sequences in peaks of height range 2-3 and higher (peak height distributions given in Fig. 2). Using the HCV sequence in plasmid Jc1FLAG2(p7-nsGluc2A) as reference (15), the ratio of transition versus transversion mutations increased in a similar proportion for the haplotypes present in low and high fitness values of HCV p100 and HCV p200. A similar increase was found for the ratio of synonymous versus non-synonymous mutations (Fig. S10A, B in https://saco.csic.es/index.php/s/7TgiQcCr9ifpnt5). Amino acid acceptability was determined with the PAM 250 matrix (44). The low acceptability substitution group (PAM 250 < 0) was less abundant in the haplotypes of the high fitness peaks of population HCV p200, but the difference with low fitness peaks was not statistically significant (Fig S10C in https://saco.csic.es/index.php/s/7TgiQcCr9ifpnt5). The comparisons of mutation types and amino acid substitution tolerance mark only tendencies in the diversification process. The fitness of the genomes whose mutations conform the haplotypes sampled in low or high fitness peaks may be dictated by mutations located anywhere in the genome. The decrease of amino acid substitutions with PAM250 < 0 in haplotypes of the high fitness peaks of HCV p200 may be due to negative selection acting on the genomes harboring them. Such a decrease is the only distinctive feature that we have identified in the mutation repertoire of the populations examined (compiled in Table S2 to S4 in https://saco.csic.es/index.php/s/7TgiQcCr9ifpnt5).

## DISCUSSION

Genetic variability of RNA (and many DNA) viruses is a major feature of their biology, and an obstacle for disease control. Numerous analyses of clinical and laboratory isolates have indicated that HCV is one of the most genetically variable RNA viral pathogens. Its plasticity results in considerable phenotypic heterogeneity that can influence disease progression and the effectiveness of antiviral interventions [reviews in (5, 6)]. Yet, the information on HCV fitness landscapes is very limited, and it has been largely restricted to genomic sequences from infected patients and centered on the effect of antiviral interventions on viral population composition. In this line, a HCV sequence database was translated into an empirical fitness landscape to design vaccines that might simultaneously decrease viral fitness and avoid selection of escape mutants (45).

Our previous studies evidenced wide and dynamic diversification of mutant spectra in the evolution from the initial HCV p0 to HCV p100 and HCV p200 (11, 13, 18); haplotype alignments are available in https://saco.csic.es/index.php/s/586L2f9jJQtbRXq). As an interpretation of the results, we proposed broadly diversifying selection as an attribute of viral quasispecies dynamics, manifested when viruses replicate in environments that do not experience external perturbations (24). A likely driver of broadly diversifying selection is the modification of mutant spectrum composition due to mutational input (11, 24, 46). The diversity indices quantified in previous studies did not inform of the relationships among the sequences present in mutant spectra. Such relationships have been approached in the present study with the ANN method SOM developed by Kohonen and colleagues (41, 42, 47). In this manner, the sequence information has yielded a fitness landscape of each HCV population. The SOM procedure has been previously used to determine the fitness landscape of HIV-1 clones and populations (30). Other applications have included the interpretation of patterns of cellular gene expression (48, 49), or the analysis of taxonomic clustering of cellular and viral RNA sequences (43). In connection with HCV, SOM clustering was used to investigate hepatocellular carcinoma (HCC) development as the basis for tumor differentiation and invasiveness, from expression levels of 12,600 genes in 50 HCC samples from patients with positive HCV serology (50). Also, Kohonen’s ANN were trained to predict undiagnosed HCV infections and infection risk (51).

The SOM analysis of HCV populations has revealed a two-layer fitness landscape. The first layer consists of multiple low fitness peaks that tend to form a broad platform, covering multiple points in sequence space. Although of limited extension, this platform was discernible in population HCV p0, implying that it was already initiated with the rounds of genome multiplication that followed the initial RNA transfection to produce HCVcc, and then the limited number of infection cycles to obtain population HCV p0 (Fig. 1A) (12). In the evolution towards HCV p100 and HCV p200, the populations maintained the same pattern, with low fitness peaks expanded towards larger areas of sequence space. The basal fitness platform is adorned with a limited number of protruding fitness peaks that resemble the standard representation of a rugged fitness landscape in the Wrightian sense (52). Interestingly, the function that relates the number of fitness peaks with peak height corresponds to a power law since it has the form y = ax^-b^ (the equations for different amplicons and viral populations are given in the legend for Fig. 2, and the confirmation of a straight line in a log-log plot of the same data is shown in Fig. S11 in https://saco.csic.es/index.php/s/7TgiQcCr9ifpnt5). A power law describes scale-free (non-random) processes in physics and biology that have some underlying dynamic event in their construction (53, 54). In the power law discovered with the HCV fitness landscape, the underlying force may be mutation, with the power law reflecting far more frequent pathways to reach the first fitness platform than high fitness peaks, with the latter requiring organized, non-random, clusters of mutations. Interestingly, the power law relationship was lost for HCV p100 and HCV p200 when the fused amplicons were used for the graphics (equations in the legend for Fig. 4). This may be due to a number of ambiguous genome positions that were generated in the amplicon fusion process (described in Materials and Methods). This point, as well as further penetration in the significance of this particular power law, require further research.

Despite the similar landscape morphology, the fitness maps of HCV p0, HCV p100 and HCV p200 show differences in peak distribution, either considering individual amplicons or a fused single 2D SOM network that recapitulates the information from the three amplicons (Figs. 2 and 4). In particular, the comparison evidences dynamics of peak movements, with striking differences between HCV p100 and HCV p200 despite the two populations having reached the same fitness value as measured by the standard growth-competition assays (13, 17). The only biochemical parameter that we identified —and that may fuel the dynamics of change from HCV p100 to HCV p200—is a 2.6-fold larger intracellular exponential growth displayed by HCV p200 relative to HCV p100 (13). However, the three NS5B amplicons did not follow the same trajectory of fitness modification. While for amplicons 1 and 3 the ratio of peaks or haplotypes unique to the population to those shared by other populations was the same for HCV p100 and HCV p200, for amplicon 2 it was two times higher for HCV p200 than HCV p100 (derived from the graphics of Figs. 2 and 4, and included in Table S1 in https://saco.csic.es/index.php/s/7TgiQcCr9ifpnt5).

The comparison of fitness landscapes has not revealed traceable evolutionary trajectories. No minority haplotypes in the ancestral HCV populations are the ancestors of most haplotypes that stand as dominant in subsequent populations. Rather, the picture obtained is that of a network of interconnected, transient sequences that do not define linear evolutionary events. Despite the absence of sub-lineages with temporal continuity, the number of identical fitness peaks that arose in independent passage replicas of the same starting population is remarkable [50.4% (range 37.5% - 72.7%) of the total for replicas (a), (b), and (c) of populations HCV p0, HCV p100, and HCV p200, subjected to four serial passages in Huh-7.5 cells (Table S5 in https://saco.csic.es/index.php/s/7TgiQcCr9ifpnt5)]. Similarity of behavior in separate evolutionary viral lineages suggests a component of determinism (predictability) in a system whose evolution should be strongly directed by stochastically arising mutations. This paradoxical behavior has been previously observed in different studies with other RNA viruses, and a number of possible underlying mechanisms have been proposed (55-59).

A more realistic perception of the complexity of the HCV fitness landscape can be obtained by considering that the SOM maps have been constructed with haplotypes from amplicons that cover only 10% of the entire HCV genome. This is a limitation of our study, although achieving a similar depth of mutation detection for whole genome amplicons than short amplicons is still technically challenging. A more populated basal platform than displayed in the SOM graphics of Figs. 2 and 4 is predicted if the analysis of haplotype frequencies were extended to additional genomic sites. The reason is that the sites of heterogeneity -defined as those with more than one nucleotide, revealed by Sanger sequencing—were found along the entire genome of the same HCV populations (11).

The fitness landscapes resulting from HCV replication in a monotonous environment can serve as a basis for comparison with the landscapes acquired when a selective constraint is applied to the evolving population. In particular, the analysis should reveal if alternative mutational pathways are available to the virus to respond to a specific constraint. Also, how the HCV fitness is shaped in patients versus the cell culture environment may be informative of adaptive mechanisms, and such a work is now in progress.

A two-layer fitness distribution may have biological consequences. The first layer or platform may prevent mutations from driving genomes into low fitness pits. It may also act as a spring-board for viral populations to reach higher fitness peaks. This should reduce the transition time between fitness peaks which is a limitation of adaptability recognized in general evolutionary genetics (60-62).

## MATERIALS AND METHODS

### Origin of the HCV populations, and serial passages in Huh-7.5 reporter cells

The initial HCVcc population was obtained by *in vitro* transcription of plasmid Jc1FLAG2(p7-nsGluc2A) (15), followed by RNA electroporation into Huh-Lunet cells, and further amplification in Huh-7.5 reporter cell monolayers to yield the parental population HCV p0 (12). HCV p0 was further passaged in Huh-7.5 reporter cells to obtain HCV p100 and HCV p200 (HCV p0 that has been propagated 100 and 200 times in Huh-7.5 reporter cells, respectively), as has been previously described (13). The sequences to derive the SOM-based fitness landscape were obtained from the three parental HCV p0, HCV p100 and HCV p200 populations subjected to further passages in two different experiments (experiment 1, and experiment 2) and several replicas. This yielded the 44 populations for which a fitness landscape was determined (populations depicted as empty circles in Fig. 1A). The sequences on which the present study is based have been previously reported (11, 18), and are available in (https://saco.csic.es/index.php/s/586L2f9jJQtbRXq). To control for the absence of cross-contamination with virus from another population or replica, mock-infected cells were maintained in parallel with each infected culture, and each supernatant was titrated; no infectivity in the mock-infected cultures was detected in any of the experiments.

Experiments of short-term evolution (up to 10 serial passages) starting from HCV p0, HCV p100 and HCV p200 (Fig. 1A) were carried out also in Huh-7.5 reporter cells. To initiate serial passages, 4 x 10^5^ Huh-7.5 reporter cells were infected with HCV p0, HCV p100 and HCV p200 at a multiplicity of infection (MOI) of 0.03 TCID_50_/cell; for subsequent passages 4 x 10^5^ fresh, Huh-7.5 reporter cells were infected with the virus contained in 0.5 ml of the cell culture medium from the previous infection of the same lineage; the multiplicity of infection (MOI) ranged from 0.1 to 0.5 TCID_50_ per cell. In all passages, infections were allowed to proceed for 72h to 96h. Additional procedures, including titration of infectivity to determine TCID_50_ values, and viral RNA quantification have been previously described (11-13).

### RNA extraction, viral RNA amplification and ultra-deep sequencing of cell culture populations

Intracellular viral RNA was extracted from the initial HCV p0, HCV p100 and HCV p200 populations, and their passaged derivatives using the Qiagen RNeasy kit (Qiagen, Valencia, Ca, USA). HCV RNA was amplified by RT-PCR using Accuscript (Agilent), and specific HCV oligonucleotide primers that have been previously described [Table S10 of (11)]. Agarose gel electrophoresis was used to analyze the amplification products, using Gene Ruler 1 Kb Plus DNA ladder (Thermo Scientific) as molar mass standard. To ascertain absence of contaminating templates, all experiments included negative controls without template RNA. To avoid sequence representation biases due to redundant amplifications of the same initial RNA templates due to template molecule limitations, amplifications were carried out with template preparations diluted 1:10, 1:100 and 1:1000; only when at least the 1:100 diluted template produced a visible DNA band, was molecular cloning performed using the DNA amplified from the undiluted template sample. PCR products were purified (QIAquick Gel Extraction Kit, QIAgen), quantified (Pico Green assay), and tested for quality (Bioanalyzer DNA 1000, Agilent Technologies) prior to Illumina deep sequencing analysis (MiSeq platform, with the 2×300-bp mode with v3 chemistry).

Several control experiments were performed in preparation of the ultra-deep sequencing procedure to ensure the reliability of the mutations derived from clean reads, for proper mutant spectrum characterization. They have been previously described (63-66), and they were as follows: first, we determined the basal error of the amplification and sequencing process, using an infectious HCV cDNA clone to perform RT-PCR, the nested PCR, and ultra-deep sequencing using Illumina MiSeq. Second, we quantified the PCR recombination frequency during the amplification steps using mixtures of *wt* and a mutant clone to perform RT-PCR, and the nested PCR and ultra-deep sequenced using Illumina MiSeq. Third, we ascertained the similarity of read composition in different RT-PCR amplifications and sequencing runs, using different samples of the same RNA preparation. We concluded that mutations identified with a frequency above the 0.5% cut-off value, and that were consistently found in the two DNA strands were considered for the analyses. For additional details of the read cleaning procedures, criteria for mutation acceptance, and experimental controls with reconstructed HCV RNA mutant mixtures, see (63-66).

### SOM derivation

Detailed description of the SOM algorithm has been published elsewhere (43). The ANN model (41) exhibits an architecture consisting of a set of neurons arranged in a rectangular grid that define a neighborhood relationship. The map size has been chosen to ensure sufficiently dispersion of the sequences mapped in the grid, while preserving the grouping of those that are similar; the resulting size is a function of the size of the data set. In the case of mapping each amplicon, a 15 x 15 size grid was selected as suitable, while for the map generated with all the fused amplicons, the selected size was 25 x 25, due to the greater number of sequences.

Every neuron has an associated prototype vector with the same nature and dimension that the input data set (in this work, the amplicon sequences). SOM generates a projection or mapping of the input space, usually high-dimensional, in the two-dimensional topological structure of the network. The SOM training algorithm determines the way in which this mapping is created. This process iteratively modifies the SOM prototype vectors to fit them to the distribution of the input data space, using a methodology similar to a regression. During the SOM training, each input vector is associated with the neuron that best matches with the pattern in any metric (the so-called ‘best matching unit’, bmu). As a result of this process, the prototype vectors associated with the bmu, and all the neurons located in a neighborhood area around it, are modified in order to move them closer to the input vector. In this work the bmu has been calculated in terms of Euclidean distance. In the case of classification of vectors with sequence data, the algorithm requires a previous transformation into equivalent numeric vectors. This has been done using the previously described codification (30, 43). Each nucleotide is transformed into the corresponding 3D numerical coordinates in an irregular tetrahedron (Fig. S1 in https://saco.csic.es/index.php/s/7TgiQcCr9ifpnt5).

In this way, each RNA sequence is transformed into a numerical vector of a dimension which is three times the length of the sequence, and this is the vector that is used by the SOM algorithm during the training process. After the training, SOM can determine similarities over the input vectors (amplicon sequences), in the sense that similar sequences will be mapped by the same neuron or by a neighboring neuron.

With the dataset of each amplicon, 25 SOM networks of size 15 x 15 were trained, and the one with the lowest Kaski-Lagus error (ε_k-l_) was selected. The same training factors were used: number of input neurons (N); dimension of the dataset vectors (length of the sequence times 3); size of the output map 15 rows times 15 columns; hexagon neighborhood connection (each neuron has six neighbors around it: two at the top, two at the bottom, one on the left, and one on the right); initial neighborhood of 14 rows and 14 columns, with neighborhood decrement at the end of each epoch (equivalent to the number of sequences in the dataset); learning factor α (t) = α1 (1-t / α2), with α1 = 0.1 and α2 equal to the total iterations of the training algorithm; the total number of iterations is equal to the total number of epochs times the number of sequences. The total of epochs is determined by the initial neighborhood + 5, that is, the algorithm carries out the necessary epochs so that the neighborhood area decreases until it affects only the bmu, plus 5 additional fine-tuning epochs (total iterations: 19 times number of dataset sequences).

Finally, a labeling process was applied to each map. Using as a basis the network selected for each dataset, the 3D fitness maps labeling was generated with the accumulated frequencies for each haplotype, so that each neuron was assigned the sum of frequencies of the haplotypes for which it is the bmu. This value represents the cumulative frequency of sequences that fall in the Voronoi region of the neuron. Although the SOM map is generated or trained with all the sequences of each dataset, the 3D maps can be obtained with the subset of sequences to be represented.

### Amplicon fusion method

Based on the fact that the amplicons have overlapping sequences (Fig. 1B), we completed for each haplotype of an amplicon a length of 1005 nucleotides using the missing information provided by the haplotypes of the other two amplicons in the same population, passage number and experiment. To achieve this for amplicon A1 (original length 312 bases), the amplicon A2 haplotypes with initial overlapping sequences matching the last 21 bases of haplotype A1 were located. The same operation was conducted with the amplicon A3 haplotypes whose initial overlapping sequences matched the last 27 bases of any of the A2 haplotypes found in the previous step. Fusion sequences were obtained for the A2 and A3 haplotype lists. To generate the final fusion sequence, the bases of each position were compared, keeping the base for any position with identical nucleotide in all sequences, or the IUPAC nucleotide ambiguity code associated with the combination of the bases when a position had more than one nucleotide. The 312 bases of the amplicon A1 haplotype were completed by adding the last 297 bases of the A2 fusion sequence followed by the last 396 bases of the A3 fusion sequence. A similar procedure was used to derive the amplicon A2 (original length of 318 bases) haplotype fusion sequence. The amplicon A1 haplotypes with final overlapping sequences matching the first 21 bases of haplotype A2, and the amplicon A3 haplotypes with initial overlapping sequences matching the last 27 bases of haplotype A2 were located. The fusion sequence was obtained for the A1 haplotype list and for the A3 haplotype list. The 318 bases of the amplicon A2 haplotype were completed including the first 291 bases of the A1 fusion sequence at the beginning and the last 396 bases of the A3 fusion sequence at the end, as described to complete amplicon 1. Likewise, for amplicon A3 (original length of 423 bases), the amplicon A2 haplotypes with final overlapping sequences matching the first 27 bases of haplotype A3, and the amplicon A1 haplotypes with final overlapping sequences matching the initial 21 bases of any of the A2 haplotypes found in the previous step were located. The fusion sequence was obtained for the A1 haplotype list and for the A2 haplotype list. The 423 bases of the amplicon A3 haplotype were completed including at the beginning the first 291 bases of the A1 fusion sequence followed by the first 291 bases of the A2 fusion sequence. When no haplotypes with matching overlapping sequences were found in any of the other two amplicons, the full list of haplotypes of the mismatched amplicon was used to equalize the length.

### Statistics

The statistical significance of differences among the number of fitness peaks and among the number of haplotypes of HCV p0, HCV p100 and HCV p200 was calculated with the t-test since the data follow a normal distribution (p > 0.05; Shapiro-Wilk test). The differences between the distribution of fitness peaks of HCV p0, HCV p100 and HCV p200 for each amplicon was calculated with the Pearson’s chi-square test. The comparison between the number of unique peaks and the number of shared peaks of HCV p0, HCV p100 and HCV p200 for each amplicon, was calculated with a proportion test. All calculations were carried out using software R version 4.0.2.

## Data availability

The Illumina data have been deposited in the NCBI BioSample database under accession numbers SAMN18645452, SAMN18645453, SAMN18645456, SAMN18645457, SAMN18645460, SAMN18645463, SAMN18645464 and SAMN18645467 (BioProject accession number PRJNA720288) for experiment 1 and SAMN13531332 to SAMN13531367 (BioProject accession number PRJNA593382) for experiment 2. The haplotypes alignments are available in https://saco.csic.es/index.php/s/586L2f9jJQtbRXq. The fasta files are termed according to the experiment (Exp1, Exp2a, Exp2b or Exp2c), population (HCV p0, HCV p100 or HCV p200), passage (initial, p1, p2, p3, p4 or p10) and amplicon (A1, A2 or A3).

## ACKNOWLEDGMENTS

The work at UPM was supported by grants TIN2017-085727-C4-3-P (DeepBio) from Ministerio de Ciencia, Innovación y Universidades (MCIU) and PID2019-104903RB-100 (Funded by the EU under the FEDER program). Work at CBMSO was supported by grants SAF2014-52400-R from Ministerio de Economía y Competitividad (MINECO), SAF2017-87846-R and BFU2017-91384-EXP from MCIU, PI18/00210 from Instituto de Salud Carlos III (ISCIII), S2013/ABI-2906 (PLATESA from Comunidad de Madrid/FEDER), and S2018/BAA-4370 (PLATESA2 from Comunidad de Madrid/FEDER). C.P. is supported by the Miguel Servet program of the Instituto de Salud Carlos III (CP14/00121 and CPII19/00001), cofinanced by the European Regional Development Fund (ERDF). CIBERehd (Centro de Investigación en Red de Enfermedades Hepáticas y Digestivas) is funded by Instituto de Salud Carlos III. Institutional grants from the Fundación Ramón Areces and Banco Santander to the CBMSO are also acknowledged. The team at CBMSO belongs to the Global Virus Network (GVN). Work at Centro Nacional de Microbiología (ISCIII) was supported by grants SAF2016-77894-R from MINECO and PI13/02269 from ISCIII. C.G.-C. is supported by predoctoral contract PRE2018-083422 from MCIU. B.M.-G. is supported by predoctoral contract PFIS FI19/00119 from ISCIII, cofinanced by Fondo Social Europeo (FSE). The work at Universidad Complutense Madrid has been supported by research grant CTQ2017-87864-C2-2-P, from the Ministerio de Economia y Competitividad (MINECO), Spain.

